# FOXO dictate initiation of B cell development and myeloid restriction in common lymphoid progenitors

**DOI:** 10.1101/2022.01.21.477216

**Authors:** Lucía Peña-Pérez, Shabnam Kharazi, Nicolai Frengen, Aleksandra Krstic, Thibault Bouderlique, Julia Hauenstein, Minghui He, Ece Somuncular, Xiaoze Li Wang, Carin Dahlberg, Charlotte Gustafsson, Ann-Sofie Johansson, Julian Walfridsson, Nadir Kadri, Petter Woll, Marcin Kierczak, Hong Qian, Lisa Westerberg, Sidinh Luc, Robert Månsson

**Affiliations:** Center for Hematology and Regenerative Medicine Huddinge, Karolinska Institutet, Stockholm, Sweden; Department of Laboratory Medicine, Karolinska Institutet, Stockholm, Sweden; Department of Medicine Huddinge, Huddinge, Karolinska Institutet, Stockholm, Sweden; Department of Microbiology Tumor and Cell Biology, Karolinska Institutet, Stockholm, Sweden; Department of Cell and Molecular Biology, National Bioinformatics Infrastructure Sweden, Science for Life Laboratory, Uppsala University, Uppsala, Sweden; Science for Life Laboratory, Department of Medicine Solna, Karolinska Institutet, and Division of Infectious Diseases, Karolinska University Hospital, Stockholm, Sweden; Hematology Center, Karolinska University Hospital, Stockholm, Sweden

## Abstract

The development of B cells relies on an intricate network of transcription factors critical for developmental progression and lineage commitment. In the B cell developmental trajectory, a temporal switch from predominant *Foxo3* to *Foxo1* expression occurs at the CLP stage. Utilizing VAV-iCre mediated conditional deletion, we found that the loss of FOXO3 impaired B cell development from LMPP down to B cell precursors, while the loss of FOXO1 impaired B cell commitment and resulted in a complete developmental block at the CD25 negative proB cell stage. Strikingly, the combined loss of FOXO1 and FOXO3 resulted in the failure to restrict the myeloid potential of CLPs and the complete loss of the B cell lineage. This is underpinned by the failure to enforce the early B-lineage gene regulatory circuitry upon a predominantly pre-established open chromatin landscape. Altogether, this demonstrates that FOXO3 and FOXO1 cooperatively govern early lineage restriction and initiation of B-lineage commitment in CLPs.

**SUMMARY:** Common lymphoid progenitors co-express the transcription factors FOXO1 and FOXO3. Removing FOXO1 and FOXO3 at this developmental stage results in regained myeloid potential, failed establishment of the early B cell gene regulatory program, and the complete loss of the B cell lineage.

## INTRODUCTION

The development of highly specialized cell types from hematopoietic stem cells depends on the expression of lineage specific transcription factors that together orchestrate the establishment of specific transcriptional programs and lineage commitment. Lymphoid development is initiated with the generation of lymphoid-primed multipotent progenitors (LMPP) (Adolfsson et al., 2005; Mansson et al., 2007) that subsequently give rise to common lymphoid progenitor (CLP) (Kondo et al., 1997). The heterogeneous CLP compartment can be divided into LY6D^-^ CLPs (CLP-A) that maintain the potential to generate all lymphoid lineages and LY6D^+^ B-lineage specified CLPs (CLP-B) (Inlay et al., 2009; Mansson et al., 2010). Within the CLP-B compartment the potential to generate NK- and T-lineage cells is restricted and B-lineage commitment occurs (Mansson et al., 2008; Jensen et al., 2018). The committed CLP-Bs differentiate into CD19 expressing B cell progenitors that undergo V(D)J rearrangement of first the Ig heavy (IgH) chain gene in proB cells and subsequently Ig light (IgL) chain genes in CD25 expressing preB cells (Rolink et al., 1994; Hardy et al., 2007; Rothenberg, 2014). The formation of a functional B cell receptor (BCR) results in the generation of immature B cells that further mature into follicular and marginal zone B cells.

The initiation of lymphopoiesis and early B cell development is critically dependent on the stage-specific expression of an interdependent network of transcription factors (Mandel and Grosschedl, 2010; Sigvardsson, 2018). At the CLP stage, early B cell factor 1 (*Ebf1*) is needed for the establishment of the early B cell transcriptional program (Lin and Grosschedl, 1995; Zandi et al., 2008) and subsequently for the activation of PAX5 (Decker et al., 2009). With EBF1 and PAX5 activated, a mutually enforcing positive feedback loop is established (Roessler et al., 2007; Decker et al., 2009; Sigvardsson, 2018) that locks down B cell identity (Nutt et al., 1999; Cobaleda et al., 2007b; Nechanitzky et al., 2013) and promotes the development of mature B cells (Gyory et al., 2012; Cobaleda et al., 2007a).

The Forkhead box O (FOXO) family of transcription factors are major effectors of the PI3K-AKT signaling pathway and out of the four family members, *Foxo1*, *Foxo3,* and *Foxo4* are expressed in the hematopoietic system. *Foxo1* is central to the B cell lineage (Szydłowski et al., 2014; Baracho et al., 2011). In a developmental context, the loss of FOXO1 was initially reported to have a mild phenotype in early B cell development {Dengler:2008ec} with no significant reductions in proB and preB cell numbers despite the role of FOXO1 in regulating the RAG genes critical for V(D)J recombination (Amin and Schlissel, 2008; Dengler et al., 2008). In addition to demonstrating that the loss of FOXO1 impairs the initiation of B cell specification in CLPs, our later study using earlier conditional deletion found a more severe reduction in B cell numbers and indicated that B cell development was blocked at the proB cell stage (Mansson et al., 2012). However, the existence of B cells in both the BM and spleen suggests that FOXO1 is not an absolute requirement for B cell commitment nor for developmental progression past the proB cell stage (Mansson et al., 2012). In addition to *Foxo1*, the loss of *Foxo3* has been shown to impact early B cell development by causing reduced preB cell numbers (Hinman et al., 2009) while a potential role in earlier lymphoid progenitors remains to be explored.

Here we utilized VAV-iCre mediated conditional deletion of FOXO1 and FOXO3 throughout the hematopoietic system to study their roles in B cell development. We show that the loss of FOXO3 impacts B lymphopoiesis at a much earlier stage than previously reported with FOXO3 being required for the generation of LMPPs, CLPs, and B cell precursors while being dispensable for the mature B cell subsets. In addition, we show without ambiguity that the loss of FOXO1 causes a developmental block at the proB cell stage mirroring that of RAG deficient mice. Strikingly, we further show that the combined loss of FOXO1 and FOXO3 resulted in a failure to restrict the myeloid potential of CLPs and to initiate the early B cell program. This resulted in the complete loss of the B cell lineage and demonstrates that cooperatively FOXO1 and FOXO3 are indispensable for the initiation of B cell development.

## RESULTS

To analyze the expression of the FOXO-family genes, we performed RNA sequencing on wild-type (WT) cells representing the B cell developmental pathway ranging from hematopoietic stem cells to mature peripheral B cells. This revealed a clear temporal switch where *Foxo3* is highly expressed in stem- and multipotent progenitor cells but downregulated in CLPs where *Foxo1* is upregulated instead and subsequently becomes the predominantly expressed FOXO gene in the B cell lineage (Fig. 1A). With both CLPs and B cells co-expressing significant levels of *Foxo3* and *Foxo1*, this prompted us to investigate more closely their role in B cell development. To this end, we utilized Vav-iCre (de Boer et al., 2003) to achieve conditional loss of FOXO1 (FOXO1ko) and FOXO3 (FOXO3ko) alone or in combination (FOXOdko) throughout the hematopoietic system. This model provided effective deletion of the loxP flanked (floxed) DNA binding domains of both *Foxo1* and *Foxo3* (Fig. S1A-C). Mice lacking the Vav-iCre transgene (mainly *Foxo1^flox/flox^Foxo3^flox/flox^* mice) were utilized as normal littermate controls (CTRL).

**Figure 1.**
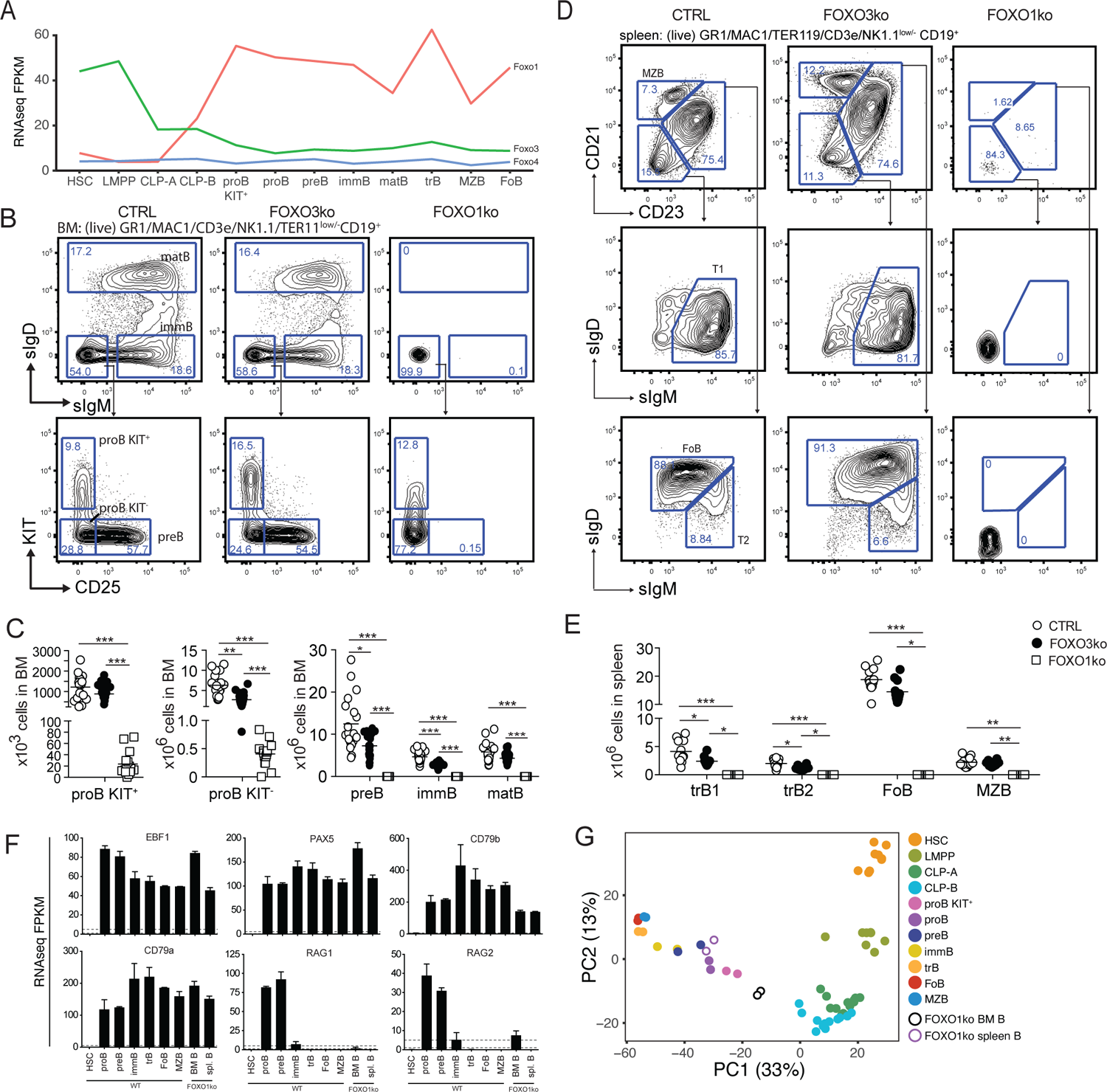
Loss of FOXO1 or FOXO3 impairs B cell development. (A) Gene expression (FPKM) of *Foxo1*, *Foxo3,* and *Foxo4*. HSC, hematopoietic stem cell; LMPP, lymphoid primed multipotent progenitor; CLP, common lymphoid progenitor; immB, immature B; matB, mature B in BM; trB, transitional B; FoB, follicular B; MZB, marginal zone B. (B) Gating strategy for identification of BM B lineage cells. (C) Total number of B lineage cells in BM. In panels C and E: each dot represents data from an individual mouse; p-values were calculated using the Kruskal Wallis test with Dunn’s test of multiple comparisons; *, ** and *** indicating p-values <0.05, <0.01, and <0.001 respectively. (D) Gating strategy for identification of spleen B lineage cells. (E) Total number of B lineage cells in spleen. (F) Expression of indicated genes. (G) Principal component analysis of RNAseq data.

### Loss of FOXO3 impairs the generation of LMPPs, CLPs, and B cell progenitors

To investigate the functional impact of the loss of FOXO3, we characterized the composition of lymphoid progenitors and B-lineage cells in BM and spleen. We found that total B cell numbers were reduced in the BM while no significant change was found in the spleen (Fig. S2A-D). Looking at the B cell progenitors in the BM of FOXO3ko mice, we found that the numbers of KIT^+^ proB cells was not reduced but a significant reduction in KIT^-^ proB cells, preB, and immature B cells was found (Fig. 1B-C). In contrast to the B cell progenitors, mature B cell numbers in the BM were not affected (Fig. 1B-C). In line with the observations from the BM, transitional B cells were reduced while follicular and marginal zone B cells were present in normal numbers in the spleen of FOXO3ko mice (Fig. 1D-E). Hence, the loss of FOXO3 impairs the generation of immature and transitional B cells from KIT^+^ proB cells while being dispensable for the maintenance of the mature B cell pool.

With *Foxo3* being highly expressed in LMPPs and CLPs (Fig. 1A), we next analyzed the progenitor compartment to investigate if FOXO3 impacted also the development of early lymphoid progenitors. While FOXO1 has been implicated in the regulation of *Il7r* expression (Kerdiles et al., 2009; Ouyang et al., 2009; Dengler et al., 2008), we found that IL7R expression readily allowed for identifying CLPs regardless of the loss of FOXO1 and FOXO3 (Fig. 2A). Analyzing the BM of FOXO3ko mice, we found that the loss of FOXO3 caused a decrease in LMPP, CLP-A, and CLP-B numbers (Fig. 2A-B).

**Figure 2.**
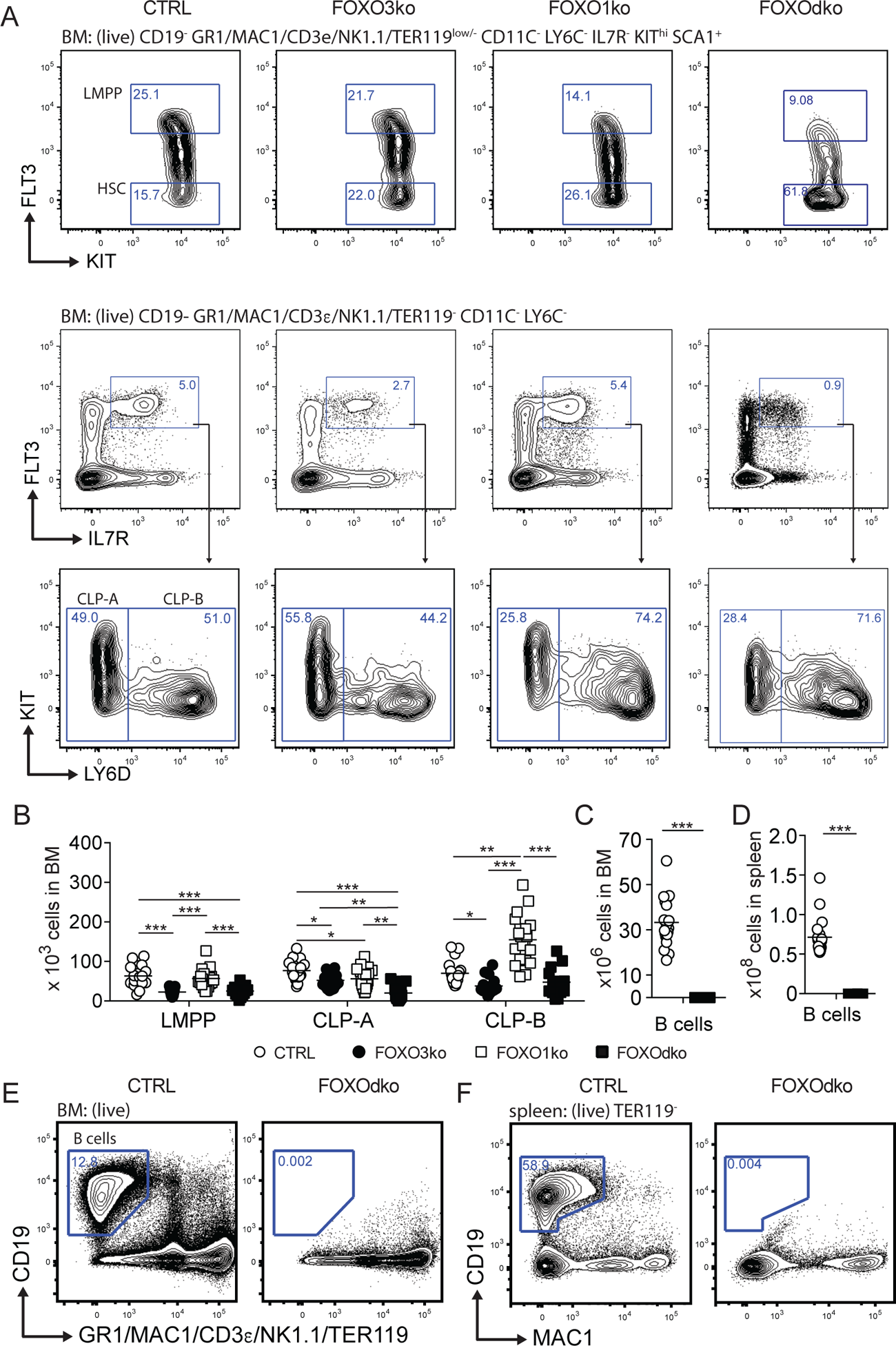
The B lineage developmental pathway is dependent on FOXO1 and FOXO3. (A) Gating strategy for identification of stem and lymphoid precursor cells. For complete gating see Fig. S3A. (B) Total number of progenitor cells in BM. (C-D) Total number B cells in (C) BM and (D) spleen. In panel B-D: each dot represents data from an individual mouse; p-values were calculated using the Kruskal Wallis test with Dunn’s test of multiple comparisons; *, ** and *** indicating p-values <0.05, <0.01, and <0.001 respectively. (E-F) Gating strategy for identification of B cells in BM and spleen.

Hence, in keeping with the *Foxo3* expression pattern (Fig. 1A), loss of FOXO3 impacts the generation of the lymphoid progenitor branch from LMPPs to transitional B cells. Altogether, this shows that FOXO3 has a previously undescribed role in the generation of lymphoid progenitors and that the role in B cell progenitors is not restricted only to the preB cell stage (Hinman et al., 2009).

### B cell development is blocked at the proB cell stage in the absence of FOXO1

In line with our earlier observations (Mansson et al., 2012), we found that the FOXO1ko mice displayed very reduced total B cell numbers in the BM (Fig. S2A and C) resulting from diminished proB cell numbers (Fig. 1B-C), a complete lack of B cell progenitors expressing CD25 (Fig. 1B-C), and a preBCR/BCR either on the cell surface (Fig. 1B-C) or intracellularly (Fig. S2E). Considering the apparent lack of cells passing beyond the proB cell stage in the BM, we next confirmed that the FOXO1ko mice had a splenic B cell population (Fig. 1D-E, S2B, and S2D) (Mansson et al., 2012). To investigate if the remaining B cells represented cells that had matured despite the lack of FOXO1, we carefully analyzed the remaining B cells. We found no expression of the conditionally deleted *Foxo1* exon (Fig. S1C), suggesting that this was not caused by escape from VAV-iCre mediated deletion. The remaining FOXO1ko splenic B cells did not generate apparent CD21/CD23-expressing populations (Fig. 2D-E) and, like the FOXO1ko BM B cells, lacked both surface (Fig 1D-E) and intracellular (Fig S2F) BCR expression. Further supporting the notion that the remaining splenic FOXO1ko B cells were not functional mature B cells, we found that the splenic architecture was not maintained (Fig. S2G) and no discernable serum levels of IgM or IgG (Fig. S2H). This indicated that the splenic B cells in the FOXO1ko represent a B cell progenitor population.

To confirm the cellular identity of the remaining B cells, we performed RNAseq on the FOXO1ko CD19^+^ cells from both BM and spleen. As expected from the CD19 expression (Fig. S2C-D), the FOXO1ko B cells maintained expression of the identity defining transcription factors *Pax5* and *Ebf1* (Nechanitzky et al., 2013; Nutt et al., 1999) as well as their target genes *Cd79a* and *Cd79b* (Sigvardsson et al., 2002; Maier et al., 2004; Akerblad et al., 1999) (Fig. 1F). Consistent with the immature cell-surface phenotype of the remaining B cells, principal component analysis (PCA) positioned the BM and spleen FOXO1ko B cells adjacent to KIT^+^ proB and total (KIT unfractionated) proB respectively (Fig. 1G). Further, in line with a developmental block in cells lacking pre-BCR expression and the known role of FOXO1 in regulating the recombinase activating genes (*Rag1* and *Rag2*) (Amin and Schlissel, 2008), we found that the expression of both Rag genes was dramatically reduced with *Rag1* expression essentially being abolished (Fig. 1F).

To establish if the loss of FOXO1 was permissive for BCR negative B cell progenitors to appear in the spleen, we next analyzed RAG2 knockout (RAG2ko) mice (Shinkai et al., 1992). Mirroring the FOXO1ko phenotype, the RAG2 deficient mice displayed a developmental block at the proB cell stage in the BM while still maintaining a population of BCR negative B cells in the spleen (Fig. S2I-K). Hence, the loss of FOXO1 is not a prerequisite for BCR negative B cell progenitors to reach the spleen.

We conclude that early loss of FOXO1, as previously suggested (Mansson et al., 2012), results in a complete developmental block at the proB cell stage. Further, we conclude that the splenic B cells in the FOXO1ko are not a sign of developmental progression but rather that FOXO1ko mice, like RAG deficient mice, harbor proB cells in the spleen.

### Loss of FOXO1 and FOXO3 blocks B cell development at the CLP-B stage

Given the co-expression of *Foxo1* and *Foxo3* at the CLP stage, we hypothesized that functional redundancy could mask a critical role of the FOXO-family in early B cell development. We found that the combined loss of FOXO1 and FOXO3 (FOXOdko) further exacerbated the loss of CLP-As observed in the single FOXO3ko while CLP-B numbers were seemingly normal (Fig. 2A-B). This indicates that the expansion of the CLP-B compartment observed after the loss of FOXO1 (Fig. 2A-B) still occurs in the FOXOdko mice. In line with this as well as the close link between the FOXO family and the regulation of proliferation, the FOXOdko mice displayed markedly increased proliferation in the CLP compartment (Fig. S3B). We found no increase in the frequency of apoptotic CLPs (Fig. S3C). With both FOXO1 and FOXO3 impacting early B cell progenitors, we next analyzed BM and spleen for B cells. Strikingly, we found that the FOXOdko mice displayed a complete loss of the B cells in BM and spleen (Fig. 2C-F). This was recapitulated upon the adoptive transfer of FOXOdko progenitor cells into normal hosts (Fig. S3D-E). Further in line with the loss of the B cell lineage, we were unable to detect serum Ig in FOXOdko animals (Fig. S2H). Hence, the combined loss of FOXO1 and FOXO3 results in a complete block in B cell differentiation at the CLP-B stage.

### CLPs lacking FOXO fail to initiate expression of the early B cell program

To investigate the gene regulatory consequences of the loss of the FOXO proteins, we performed RNAseq on FACS sorted progenitor cells from the FOXO knock-out animals, WTs, and CTRLs. Using PCA, we found that the overall expression profiles of the progenitor populations from the knock-out animals corresponded well to that of their normal counterparts (Fig. 3A). Looking only at the progenitors, the populations similarly organized in developmental order (PC1) (Fig. 3B). However, the second component (PC2) clearly separated the FOXOdko cells from the other genotypes, suggesting that the combined loss of FOXO1 and FOXO3 is needed to cause a significant impact on the overall transcriptional program across the progenitor hierarchy. With B cell development being blocked at the CLP stage in the FOXOdko, we utilized EdgeR to identify significant expression changes occurring in CLP-A and CLP-B. This identified 319 genes differentially expressed in FOXOdko CLPs (Fig. 3C-D and Table S1). A substantial number of these genes overlapped with genes significantly up-regulated in the normal CLP-A to CLP-B transition (Fig. 3C and S4A). Overall, the loss of FOXO3 gave milder gene expression changes than the loss of FOXO1 and these changes were exacerbated by the combined loss of both FOXO proteins (Fig. 3E). This suggests that the FOXO-proteins cooperatively regulate these genes. The usage of gene ontology analysis on the different gene clusters (Fig. 3D), revealed that these were associated with leukocyte biology (cluster I); B cell activation and primary immunodeficiency (cluster II-III); and response to oxidative stress (cluster IV) (Tothova et al., 2007) (Fig. S4B). Looking at specific genes, we found that the loss of the FOXO genes impaired the expression of genes critical for the B cell lineage including the surrogate light chain genes (*Igll1*, *VpreB1,* and *Vpreb3*), signaling molecules (*Cd79a*, *Blk,* and *Blnk*), and transcription factors (*Pou2af1*, *Irf4,* and *Ets1*) (Fig. 3D and F). Further, in agreement with both disruption of the positive Ebf1-Foxo1 feedback loop (Mansson et al., 2012) and the loss of the B cell lineage, *Ebf1* and *Foxo1* expression was dramatically reduced while *Pax5* expression was lost (Fig. 3D and F). In conclusion, the combined loss of FOXO1 and FOXO3 results in a failure to initiate the B cell program in CLPs.

**Figure 3.**
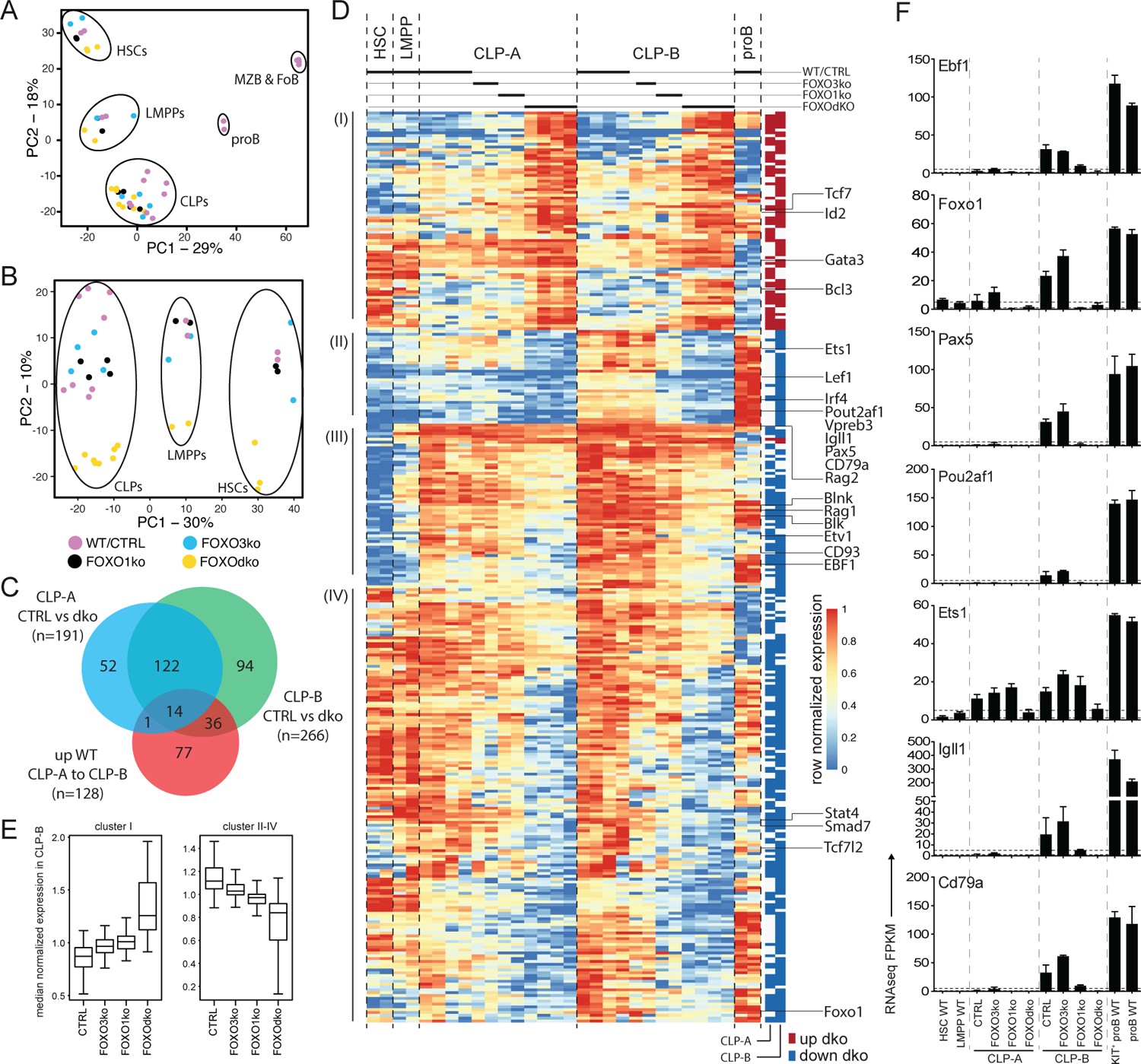
FOXOdko CLPs maintain cellular identity but fails to up-regulate B lineage related genes. (A-B) Principal component analysis of RNAseq data. (C) Venn diagram displaying the overlap between genes differentially expressed (Bonferroni corrected p-value <0.05 and 2-fold difference comparing FOXOdko to CTRL and ≥30 reads in at least two samples) in the FOXOdko at the indicated CLP stage or in the CLP-A to CLP-B developmental transition. (D) Hierarchical clustering of expression for genes displaying differential expression in FOXOdko CLPs. The stage where a significant change occurs is indicated to the right. (E) Median normalized expression of genes within clusters I and cluster II-IV from panel D. (F) Expression of indicated genes.

### FOXO is required for the establishment of the B-lineage gene regulatory landscape

To understand the gene regulatory mechanisms that cause the transcriptional changes observed in the FOXOdko CLPs, we analyzed chromatin accessibility using ATACseq (Buenrostro et al., 2013). This identified approximately 40 000 open chromatin regions (ATACseq peaks) per investigated population, with the majority of the peaks being localized distal to known promoters (Fig. S4C). Subjecting the ATACseq data to PCA, we found that the progenitor subsets organized together and in a developmental trajectory regardless of the loss of FOXO activity (Fig. 4A, left). Similar to the PCA of RNAseq data, the effect of the combined loss of FOXO1 and FOXO3 was observable when looking specifically at the progenitors (PC2) (Fig. 4A, right). Next, we used EdgeR to identify regions with highly significant changes in chromatin accessibility in the FOXOdko CLPs. This identified 297 and 1068 differentially accessible regions (DARs) at the CLP-A and CLP-B developmental stages, respectively (Fig. 4B and Table S2). The majority of the DARs were localized outside of promoter regions (Fig. 4C).

**Figure 4.**
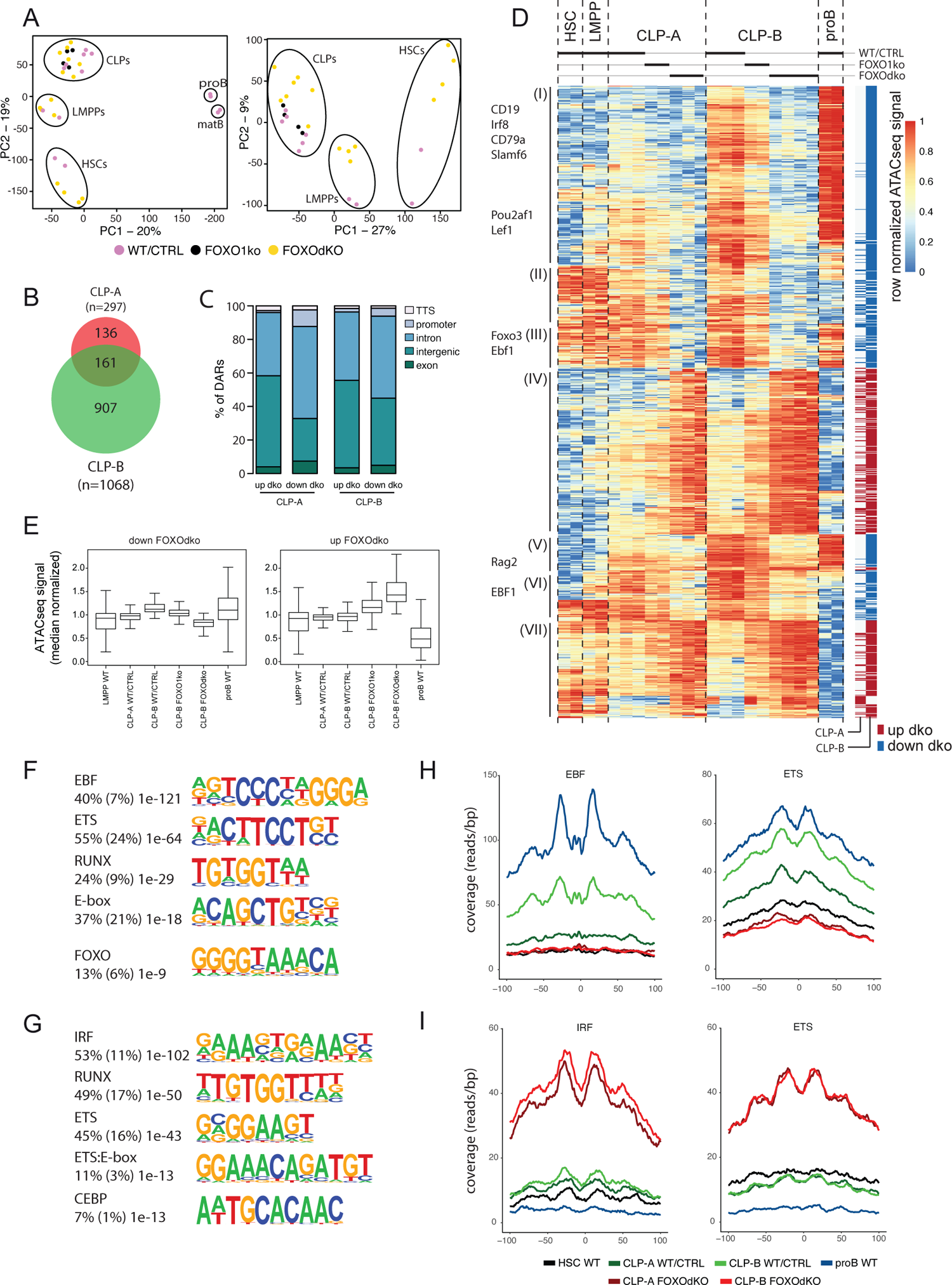
Chromatin accessibility is altered by the lack of FOXO. (A) Principal component analysis of ATACseq data. (B) Venn diagram displaying the overlap between differentially accessible regions (DARs) (adjusted p-value <0.01 and 2-fold change comparing FOXOdko to WT/CTRL and ≥30 reads in at least two samples) in the FOXOdko at the indicated CLP stage. (C) Bar graph illustrating distribution of DARs amongst indicated genomic locations. (D) Hierarchical clustering of ATACseq signal in DARs. The stage where a significant change occurs is indicated to the right. DAR proximal genes are indicated to the left. (E) Median normalized ATACseq signal in DARs with decreased (left) and increased (right). (F-G) De novo motif enrichment in DARs with decreased (F) and increased (G) ATACseq signal in FOXOdko CLP-B. Percentage of sequences containing the motif, percentage of background sequences containing the motif (in parenthesis) and p-value of the enrichment are displayed to the left of the enriched motifs. (H-I) Transposase integration-based cut-profiles of DARs with: (H) decreased ATACseq signal containing EBF and ETS binding sites; or (I) increased ATACseq signals containing IRF and ETS binding sites.

Looking at the chromatin accessibility of the DARs specifically in a B-lineage developmental context, the DARs with increased accessibility in the FOXOdko CLPs were generally regions that would lose accessibility upon developmental progression to the proB cell stage (Fig. 4D, cluster IV and VII; and 4E, right). In contrast, DARs with reduced accessibility in the FOXOdko CLPs generally displayed increased accessibility in proB cells and were localized in proximity with B-lineage associated genes including *Cd19*, *Cd79a*, *Pou2af1*, *Rag,* and *Ebf1* (Fig. 4D, cluster I, III and V; and 4E, left). This suggests that the affected elements are a critical part of the early B cell gene regulatory circuitry activated at the CLP stage.

To identify transcription factors (TFs) responsible for the changes in chromatin accessibility in the FOXOdko CLPs, we performed motif enrichment analysis on the DARs. The DARs with reduced accessibility in the FOXOdko CLPs displayed enrichment of transcription factor binding sites (TFBS) of TF families central to the B-lineage gene regulatory program including EBF, ETS, RUNX, E-box (E-protein), and FOXO (Fig. 4F). To further support the loss of binding of these TFs to the DARs with reduced accessibility, we analyzed the transposase integration in the areas surrounding the TFBS to produce cut-profiles. In line with the loss of FOXO and EBF1 binding to the DARs with reduced accessibility, we found that the cut-profiles of these TFs were reduced or completely lost (Fig. 4H and S4D). Similarly, the ETS footprint was clearly diminished (Fig. 4H) and several ETS-family genes (including *Ets1*, *Elk3,* and *Etv1*) were downregulated in the FOXOdko CLPs (Fig. 3D and S4E). We found no significant differences in the expression of the Runx-family and E-proteins, nor clear cut-profiles in the regions surrounding the TFBS of these TFs (Fig. S4D-E). Together, this suggests that the failure to express the early B-lineage program in the FOXOdko is caused by a combination of the loss of FOXO activity and the failure to properly up-regulate EBF1 and ETS-family genes.

Conversely, we found that the DARs with increased accessibility were associated with IRF, RUNX, ETS, ETS:E-box, and CEBP binding sites (Fig. 4G). No corresponding up-regulation of genes within the respective transcription factor families was observed (Fig. S4E). However, the IRF and ETS binding sites were associated with clear cut-profiles consistent with the binding of these TFs to the DARs with increased accessibility (Fig. 4I). The other TFBS did not display clear cut-profiles (Fig. S4D). Hence, this indicates that the gain of chromatin accessibility in the FOXOdko CLPs is related to the altered binding of TFs commonly expressed in CLPs.

### Open chromatin is largely established independent of FOXO in CLPs

Both FOXO1 and EBF1 have been shown to be pioneer factors (Treiber et al., 2010; Hatta and Cirillo, 2007) that can remodel condensed chromatin and make it accessible for other TFs. Given the co-dependence of *Foxo1* and *Ebf1* expression in CLPs (Mansson et al., 2012) and the essential loss of both *Foxo1* and *Ebf1* expression in the FOXOdko (Fig. 3F), we next investigated if this was associated with a failure to establish chromatin accessibility at these loci. Somewhat surprisingly, we found that the chromatin accessibility landscape surrounding both the *Foxo1* and *Ebf1* genes was essentially maintained in the FOXOdko CLPs (Fig. 5A-B). While a few DARs with decreased chromatin accessibility were identified (±1Mbp of the respective promoter regions), accessibility was not lost in these regions (Fig. 5A-B). Hence, the activation of the FOXO-EBF1 co-regulatory loop occurs via FOXO and EBF1 binding to pre-established open chromatin.

**Figure 5.**
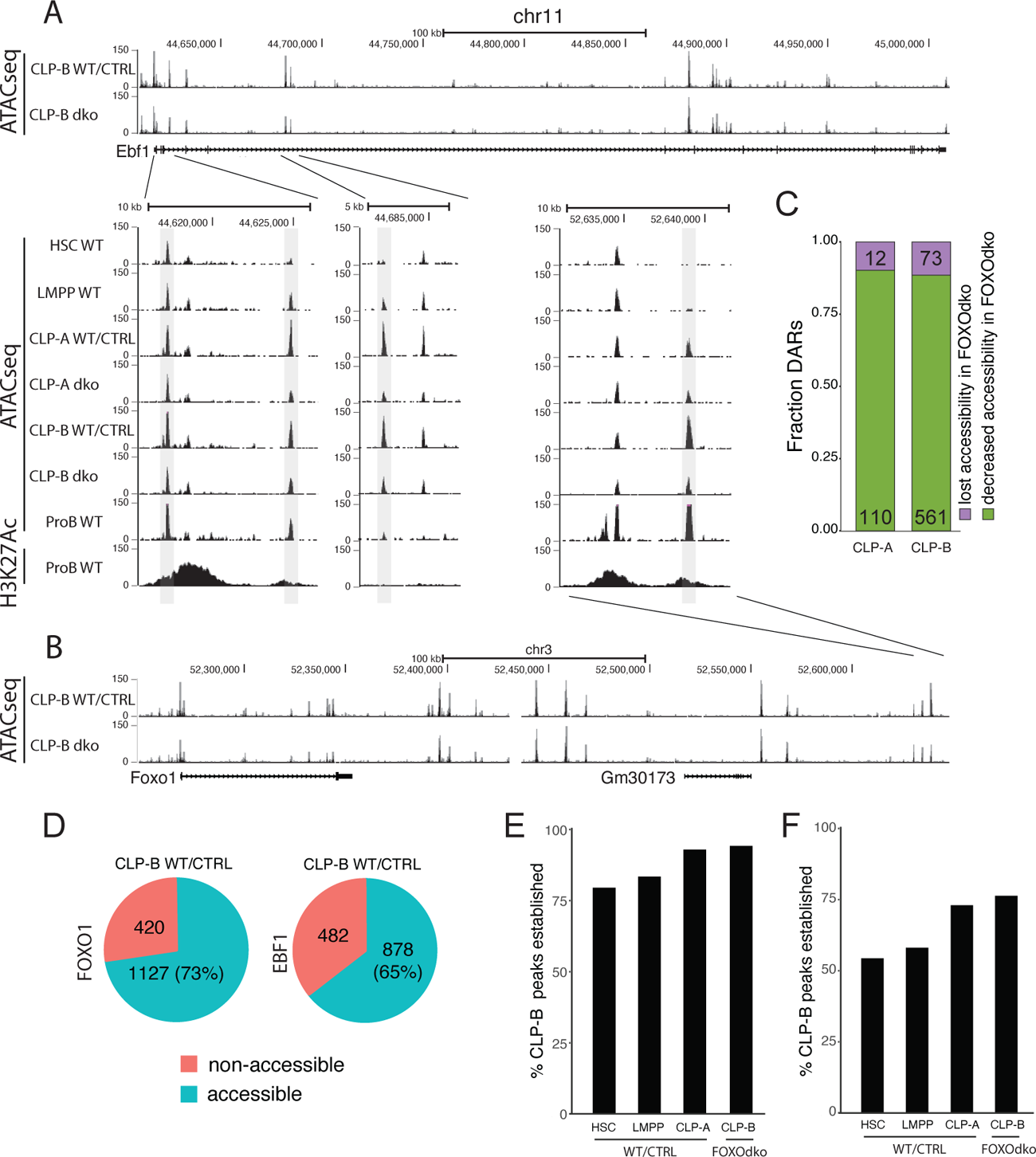
Loss of FOXO is not associated with a failure to establish open chromatin. (A-B) Tracks showing chromatin accessibility (ATACseq) and H3K27Ac (ChIPseq) for indicated cell- and genotypes at the (A) *Ebf1* and (B) *Foxo1* loci. (C) Fraction of DARs with reduced accessibility displaying either lost or decreased chromatin accessibility in the FOXOdko CLPs. (D) Venn diagram showing the overlap between FOXO1 and EBF1 binding in RAG1ko proB cells and open chromatin in WT/CTRL CLP-B. (E-F) Percentage of the open chromatin regions found in WT/CTRL CLP-B that overlapping with (E) FOXO1 or (F) EBF1 binding and that constitute open chromatin at the indicated stages of development.

To address if a similar pattern was found on a broader level, we addressed if DARs with significant decreased chromatin accessibility (Fig. 4D) were associated with a failure to establish open chromatin in the FOXOdko CLPs. To this end, we analyzed the extent to which DARs remained identified as open chromatin (peaks) in ATACseq from FOXOdko CLPs. We found that approximately 90% of the affected regions were identified as open chromatin (peaks) in FOXOdko CLPs (Fig. 5C). Thus, the reductions in chromatin accessibility observed in the FOXOdko CLPs was generally associated with decreased but not lost chromatin accessibility. To look specifically at FOXO1 and EBF1 bound regions, we overlapped FOXO1 and EBF1 identified by ChIPseq in RAG1ko proB cells (Lin et al., 2010) with open chromatin in stem- and progenitor cells. We found that 73% of FOXO1 and 65% of EBF1 bound regions constituted open chromatin (peaks) at the CLP-B stage (Fig. 5D). Out of the FOXO bound regions open in WT CLP-Bs, we found that >75% of these regions were open chromatin already in HSCs and LMPPs while >90% of the regions were open chromatin in CLP-A and FOXOdko CLP-Bs (Fig. 5E). Similarly, analyzing the EBF1 bound regions, approximately 75% of the regions were accessible (identified as peaks) in the very low *Ebf1* expressing CLP-A and FOXOdko CLP-B (Fig. 5F). In addition, just over 50% of the regions constituted accessible chromatin already in LMPPs and HSCs stages prior to the expression of *Ebf1* (Fig. 5F).

Together, this suggests that the open chromatin at gene regulatory elements needed for activation of the early B cell transcriptional program at the CLP stage is, to a large extent, pre-established by FOXO and EBF1 independent mechanisms.

### CLPs devoid of FOXO display increased ability to generate myeloid cells

Given the altered expression of genes controlling B-lineage commitment and lineage restriction in the FOXOdko, we next analyze the potential of the FOXOdko CLPs to generate T- and myeloid-lineage cells. Using the OP9-DL1 system to evaluate the potential of single progenitor cells to generate T cells, we found that essentially all growing colonies contained T cell progeny (Fig. 6A). This indicates that the ability to generate T cells is maintained in FOXOdko lymphoid progenitors. In agreement with this, analysis of the thymus revealed normal or slightly elevated numbers of γδ-, CD4, and CD8 T cells in the FOXOdko mice (Fig. S5A-B).

**Figure 6.**
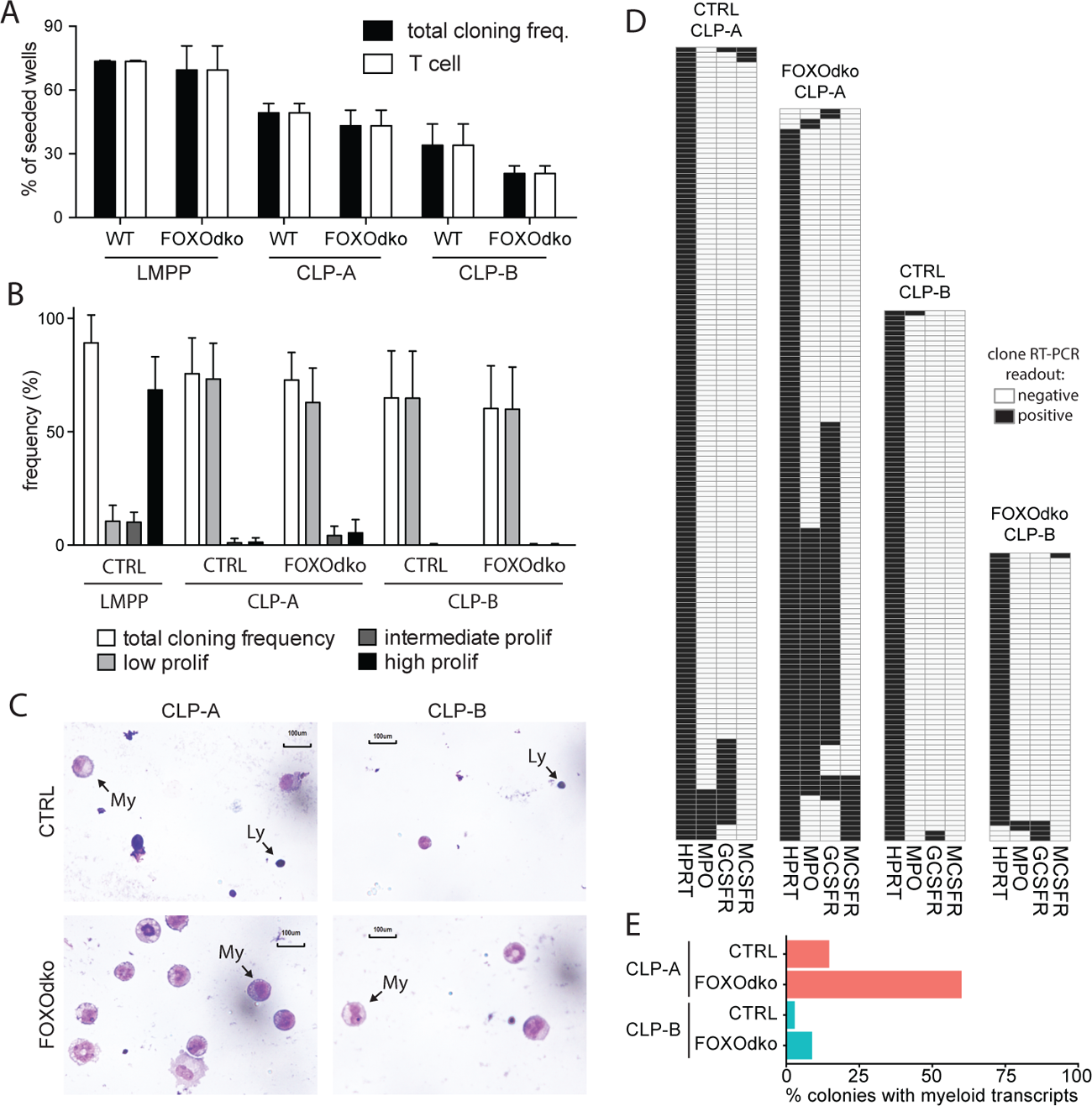
Loss of FOXO alleviates myeloid restriction in CLPs. (A) Frequency of single progenitors generating clones and T-cells on OP9-DL1. (B) Frequency of single CLPs capable of generating colonies of the indicated size under myeloid growth conditions. Data is from seven independent experiments. (C) Morphology of cells derived from CLPs under myeloid conditions. Arrows indicate lymphoid (Ly) and myeloid (My) cells. (D) Heatmaps of gene expression in single-CLP derived colonies detected by multiplex RT-PCR. Data is from three independent experiments. I Percentage of colonies expressing myeloid transcripts.

Subjecting CLPs to myeloid growth conditions, we found that the CLPs regardless of genotype gave rise to predominantly small (low proliferative) colonies with similar frequency (Fig 6B). To investigate the cellular composition of the colonies, we generated cytospins from pools of clones. As expected, CTRL CLPs mainly generated lymphocytes under myeloid conditions though limited numbers of myeloid cells could be observed, in particular in cytospins generated from CLP-A derived colonies (Fig 6C). Strikingly, we observed more abundant myeloid cells in cytospins generated from FOXOdko CLP colonies (Fig. 6C). As we were unable to generate cytospins from individual colonies because of the low number of cells per colony, we instead performed RT-PCR on individual colonies to investigate the presence of myeloid restricted transcripts on the clonal level. Consistent with the increased generation of myeloid cells, we found that an increased frequency of colonies derived from FOXOdko CLP-A expressed myeloid transcripts (Fig. 6D-E). Hence, the loss of FOXO activity results in the failure to restrict myeloid potential with FOXOdko CLP-As, on a clonal level, displaying a higher propensity to generate myeloid cells. We did not find that this was associated with significant stage specific up-regulation of TFs including *Spi1* (*Pu.1*) and the *Cebp*-family (Fig. S5C) nor of myeloid growth factor receptors commonly associated with myeloid differentiation (Fig. S5D). Interestingly, the expression of G-CSFR was down-regulated normally in the FOXOdko CLP-Bs (Fig. S5D). Together, this demonstrates that the loss of FOXO activity results in a failure to properly restrict the myeloid potential of CLP-A.

## DISCUSSION

The B cell developmental trajectory is critically dependent on TFs for developmental progression, establishing the B cell transcriptional program, and achieving B-lineage commitment. Analyzing the expression pattern of the FOXO TF family genes, we found that *Foxo3* was the dominantly expressed FOXO gene at the HSC and LMPP stages. Subsequently, the simultaneous downregulation of *Foxo3* and up-regulation of *Foxo1* at the CLP stage left *Foxo1* as the dominantly expressed FOXO-family gene in B-lineage cells. This temporal switch from *Foxo3* to *Foxo1* expression at the CLP stage together with the co-expression of *Foxo3* and *Foxo1*, in particular at the CLP-B stage, prompted us to investigate how the loss of FOXO3 and/or FOXO1 from the HSC onwards impacts the B cell developmental pathway.

In line with *Foxo3* being the dominantly expressed FOXO gene in early hematopoiesis, we found that the loss of FOXO3 significantly impaired the generation of LMPPs and CLPs adding FOXO3 to the list of TFs affecting early lymphopoiesis. While the loss of FOXO3 in the LMPPs had not previously been investigated, we also observed much broader effects on to the early B cell developmental pathway compared to prior studies (Dengler et al., 2008; Hinman et al., 2009) with essentially all steps from proB to immature and transitional B cells being affected. Potentially, this can be explained by the use of KIT, CD25, IgM, and IgD allowing us to better resolve the subtle changes to early B cell development.

As expected from *Foxo1* being upregulated at the CLP stage and in line with our previous report (Mansson et al., 2012), we found that the loss of FOXO1 caused no discernable effect on LMPP generation while CLP numbers were perturbed. Our earlier study suggested that FOXO1ko mice displayed a more severe phenotype than initially reported (Dengler et al., 2008) with a complete block at the proB cells stage (Mansson et al., 2012). However, the identity of the residual B cells remained unverified on a molecular level and it remained unresolved if splenic B cells in the FOXO1ko represented mature cells that had developed in the absence of FOXO1. Here unambiguously show that the remaining FOXO1ko B cells represent proB cells and that the FOXO1ko mice, similar to RAG deficient mice, harbor proB also in the spleen. Hence, we show here FOXO1 is absolutely required for B cell development to progress past the CD25 negative proB cell stage and that likely this is directly caused by the failure of the residual proB cells to properly express the *Rag* genes (Amin and Schlissel, 2008). The difference between the Vav-iCre and Mb1-Cre models (Dengler et al., 2008), likely is a reflection of the Mb1-Cre model (Hobeika et al., 2006) allowing B cell progenitors to proceed past early requirements for FOXO1 and EBF1 activity as the expression of *Mb1* (*Cd79a*) depends on EBF1 (Maier et al., 2004; Zandi et al., 2008).

Interestingly, we found that the FOXO1ko proB cells maintained normal *Ebf1* expression. This suggests that the FOXO1-EBF1 positive feedback loop, critical for expression of *Ebf1* at the CLP stage (Mansson et al., 2012), is less important to maintain *Ebf1* expression after B-lineage commitment and the development of proB cells. Potentially the establishment of the positive EBF1-PAX5 feedforward loop (Decker et al., 2009; Roessler et al., 2007) supersedes FOXO1-EBF1 regulation or simply decrease the need for FOXO activity to a level where FOXO3 can compensate for the loss of FOXO1 to maintain *Ebf1* expression.

The CLP-B stage is the center of the temporal switch from *Foxo3* to *Foxo1* with the genes displaying similar expression on the mRNA level. This suggested that both could act to support B cell development and that compensation by FOXO3 allowed for developmental progression and B-lineage commitment to occur in the FOXO1ko mice. As hypothesized, the combined loss of FOXO1 and FOXO3 resulted in a complete block of B cell development at the CLP-B stage. The loss of the B cell lineage was associated with the failure to initiate the early B-lineage gene regulatory program on both the transcriptional and gene regulatory level. The more exacerbated changes observed in the FOXOdko CLPs, as compared to the loss of FOXO1 or FOXO3 alone, clearly show that FOXO1 and FOXO3 synergistically activate the B-lineage gene regulatory circuitry in CLPs. This effect is clearly observable on the reduced or lost expression of several TFs critical for the B-lineage in the FOXOdko CLPs, including *Ebf1, Pax5*, *Pou2af1,* and *Ets1* (Lin and Grosschedl, 1995; Nutt et al., 1999; Schubart et al., 1996; Eyquem et al., 2004). Taken together, this suggests that the loss of the B cell lineage in the FOXOdko mice is caused by the failure to establish the coordinated expression of several transcription factors which ultimately results in a failure to express *Pax5* and undergo B-lineage commitment.

Both FOXO1 and EBF1 have been implicated to be ‘pioneer factors’ (Treiber et al., 2010; Hatta and Cirillo, 2007) that can open condensed chromatin and make it accessible for DNA binding proteins. While the pioneering function of EBF1 has been studied in the context of B cell development (Treiber et al., 2010; Li et al., 2018; Wang et al., 2020; Strid et al., 2021), it remains to be investigated if the FOXO family has a similar function. With ATACseq, in essence, defining open chromatin through the enrichment of reads originating from accessible regions, we utilized this to characterize how the loss of FOXO and EBF1 in the FOXOdko CLPs affected the establishment of open chromatin. Somewhat surprisingly, we found that despite the close association between regions with highly significant reductions in chromatin accessibility in the FOXOdko CLPs and the binding sites of FOXO and EBF1, only a minor fraction of these regions did not remain open chromatin in the FOXOdko CLPs. In addition, the majority of both FOXO1 and EBF1 bound regions established as open chromatin at the CLP-B stage were already open chromatin in HSCs. While a role of the low expressed *Foxo4* or residual Ebf1 expression for establishing open chromatin cannot be excluded, our data strongly suggests that both FOXO and EBF1 act mainly on pre-established open chromatin at the level of the CLP but that their binding causes further increased accessibility.

While the suppression of myeloid potential in committed B cells relies on PAX5, it remains unclear how the potential to give rise to myeloid cells is restricted in the LMPP to CLP developmental transition (Rothenberg, 2014). Subjecting CLPs to myeloid growth conditions, we found that the CLP-A from the FOXOdko on the clonal level displayed substantially expanded ability to generate clones containing myeloid cells compared to CLP-A with normal FOXO function. While we did observe that gains in chromatin accessibility were associated with CEBP-family binding sites in the FOXOdko CLPs, we did not find significant changes to the expression of transcription factors (including *Spi1* (*Pu.1*), *Cebpa,* and *Cebpb*) known to influence myeloid fate determination (DeKoter and Singh, 2000; Xie et al., 2004) nor of myeloid growth factor receptors. This could potentially suggest that the regulatory changes that occur are simply too subtle for us to detect them or that the loss of FOXO activity alters the response of the CLPs to the myeloid growth conditions rather than directly influencing the expression of genes limiting the myeloid potential. While a role of EBF1 in restricting the myeloid potential in CLP-As can’t be excluded, the already very low expression of Ebf1 in normal CLP-As would argue against it playing a major role.

Interestingly, we found that the G-CSFR mRNA expression was sharply downregulated in the CLP-A to CLP-B transition in both normal and FOXOdko mice. Likely, in a similar manner to the PAX5 mediated down-regulation of M-CSFR in proB cells (Nutt et al., 1999; Tagoh et al., 2006), this acts to restrict the response to myeloid growth conditions of CLP-Bs and at least in part could explain the maintained myeloid restriction in FOXOdko CLP-Bs. As this occurs in the FOXOdko CLP-Bs, which essentially lack FOXO activity as well as expression of *Ebf1* and *Pax5*, this suggests that the gene regulatory circuitry mediating myeloid restriction at the CLP stage is independent of major transcription factors known to drive B-lineage commitment.

In sum, we show that both FOXO1 and FOXO3 are needed for the proper development of lymphoid progenitors and B-lineage cells. Strikingly, the combined loss of FOXO1 and FOXO3 in CLPs results in incomplete myeloid restriction, the failure to establish the early B cell gene regulatory program, and the complete loss of the B cell lineage. Future studies aimed at identifying additional factors responsible for establishing gene regulatory features and myeloid restriction at the level of the CLP will be paramount to understanding the early lymphoid specification and lineage restriction events.

## MATERIALS AND METHODS

### Mice

To generate mice lacking FOXO1 and/or FOXO3 throughout the hematopoietic system, Vav-iCre (de Boer et al., 2003) was utilized in combination with conditional (floxed) *Foxo1* (Paik et al., 2007) and *Foxo3* (Castrillon et al., 2003) alleles. In addition, RAG2 knockout (ko) (Shinkai et al., 1992) and wildtype (WT) C57BL/6 mice were used. Genetically modified strains were maintained on a C57BL/6 background. Mice were analyzed at an age of 8-14 weeks. All animal experiments were approved by the local animal ethical committee.

### Cell preparation, flow cytometry and cell sorting

BM, spleen and thymus were dissected, crushed in PBS with 2% FCS and passed through a 70µm filter. Cells were counted using a Sysmex hematology analyzer (Sysmex). Cells were incubated with Fc-blocking antibodies (CD16/32, clone: 93, BD) prior to staining with fluorescently labeled antibodies. For antibody panels see Table S3. Dead cell discrimination was performed using propidium iodide (PI), 7AAD or live/dead fixable Aqua dead cell stain (Thermo Fisher Scientific). For FACS sorting of progenitors from BM, mature cells were depleted using purified antibodies against TER119, CD19, CD3, GR1 and MAC1 in combination with sheep anti-rat IgG Dynabeads (Thermo Fisher Scientific) prior to the specific antibody staining. For FACS sorting of B cell progenitors, total BM cells were either depleted as described above (omitting CD19) or enriched using anti-B220 beads (Miltenyi) prior to the specific antibody staining. Flow cytometry and cell sorting were performed using mainly the LSRFortessa and FACSARIAIIu/III platforms (BD Biosciences). Analysis of FACS data was done using FlowJo (BD Biosciences) and statistical analysis performed in R.

### Transplantation

Transplantation was performed by tail-vein injection of lethally irradiated 10-16 weeks old CD45.1 (B6.SJL) mice. The full irradiation dose (1000cGy) was given as two split doses. Each transplanted mouse was given 5×10^6^ BM cells from control or FOXOdko mice (CD45.2, donor cells) and 0.2×10^6^ BM cells from B6.SJL mice (CD45.1, support cells). Analysis of reconstitution in peripheral blood (PB) and spleen was performed 12 weeks post-transplantation using FACS. For antibody panels see Table S3.

### Cell cycle and apoptosis analysis

For cell cycle analysis, fully surface stained cells (stained as described above, for staining panel see Table S3) were fixed with 1% formaldehyde (Thermo Fisher Scientific), permeabilized with 0.05% Triton X-100 (Sigma) and subsequently stained with Ki67 AlexaFluor488 (BD Biosciences) and DAPI (Thermo Fisher Scientific). For analysis of apoptosis, cells were surface stained (for antibody panel see Table S3) and subsequently stained with Annexin V (Biolegend).

### ELISA

Serum was collected by tail bleeding and kept at −80°C until use. To perform the ELISA assays, MaxiSorp ELISA plates (Thermo Fisher Scientific) were coated with 10-20µg/ml of anti-mouse immunoglobulin (Ig) (H+L) (Southern Biotech) to capture total Ig in the serum. Serum Ig was detected using a secondary HRP-conjugated anti-mouse IgM (Southern Biotech) or IgG (Southern Biotech) antibody. Phosphatase substrate (Merck) was added to generate a signal subsequently measured by an ELISA plate reader.

### Immunohistochemistry

Cryostat sections (10 μm) were fixed in ice-cold acetone and then stained in phosphate-buffer saline with 5% fetal calf serum (FCS) and 0.3% Tween. The following reagents were used: B220 (RA3-6B2), TCRβ (H57-597, BioLegend); CD169 (MOMA-1); MARCO (EPR22944, AbCam), biotinylated peanut agglutinin (Vector Laboratories); goat anti-rat-AlexaFluor488, Strepavidin-AlexaFluor555 (Thermo Fisher Scientific). Images were collected with a Leica DM IRBE confocal laser scanning microscope (Leica Microsystems) equipped with 1 argon and 2 HeNe lasers, using an HC PL APO lens at 10x/0.40 CS and 90% glycerol (MP Biomedicals) and processed with Adobe Photoshop CS5 (Adobe).

### RNAseq

3-10k cells were FACS sorted into buffer RLT with β-mercaptoethanol and total RNA prepared using RNAeasy Micro (Qiagen) with on column DNase I treatment. Stranded RNAseq libraries were prepared using the TotalScript RNAseq kit (Epicenter). Libraries were sequenced paired-end (2×50 cycles) on the Illumina platform (Illumina). Reads were mapped using STAR (v 2.5.2b) (Dobin et al., 2013), strand-specific reads in exons quantified using HOMER (Heinz et al., 2010) and significant changes identified using EdgeR (Robinson et al., 2010). Only genes with ≥30 reads in at least two samples were considered in the analysis. Data was log transformed and quantile normalized for display. PCA and visualization were performed using R (v3.3.3). GO term analysis was done using Metascape (Zhou et al., 2019) with standard settings.

### ATACseq

3-10k cells were FACS sorted into cold 200μl PBS with 2% FCS and ATACseq libraries immediately prepared as previously described (Buenrostro et al., 2013). Libraries were sequenced paired-end (2×50 cycles) on the Illumina platform (Illumina). To obtain differential ATACseq peaks, data was first trimmed and mapped to mm10 using bowtie2 (v2.3.3.1) {Langmead2012}. Elimination of PCR duplicates, identification of peaks, annotation of peaks, quantification of reads in peaks and motif enrichment analysis was done using HOMER. Identification of significant changes to chromatin accessibility was done using EdgeR considering peaks with ≥30 reads in at least two samples. For visualization of data in heatmaps and box plots, data was log transformed and quantile normalized. Tracks were prepared for visualization in the UCSC genome browser (Kent et al., 2002) by calculating the median IP efficiency-normalized signal of biological replicas. Cut-profiles were made using the HOMER function annotatePeaks.pl with parameters -fragLength 9 -hist 1 inputting regions with differential accessibility (as identified by EdgeR) and containing transcription factor binding sites (TFBS) of any expressed TF (FPKM ≥ 1 in ≥2 samples) from the analyzed TF family. The cut-profiles obtained for each sample were subsequently normalized to the IP efficiency and the average across biological replicates plotted.

### ChIPseq

ChIPseq was performed as previously described (Bouderlique et al., 2019) using H3K27Ac antibodies (Diagenode Cat #C15410196, lot A1723-0041D/2) and the ThruPLEX DNA-seq (Rubicon Genomics) library preparation kit. Reads were mapped using bowtie2. Elimination of PCR duplicates and generation of tracks was performed using HOMER. Visualization performed using the UCSC genome browser.

### In vitro evaluation of myeloid potential and colony RT-PCRs

To evaluate myeloid potential of progenitors, 150 cells were FACS sorted into 3ml of medium and single cells manually plated (20µl/well) in Terasaki plates. Medium constituted IMDM (with GlutaMax, Thermo Fisher Scientific) supplemented with 100U/ml penicillin-streptomycin (Hyclone), 100µM β−mercaptoethanol, 20% Fetal Calf Serum (Merck), 25ng/ml murine KL, 25ng/ml murine GM-CSF, 25ng/ml human TPO, 25ng/ml human G-CSF, 25ng/ml human FL and 10ng/ml murine IL3 with or without 25ng/ml murine M-CSF (all from PeproTech). No apparent difference was observed between cultures with or without M-CSF and data was combined. Size of CLP- and LMPP-derived colonies was scored as low (colonies containing 2-99 cells), intermediate (≥100 cells to cells covering <10% of the well) or high (cells covered ≥10%-100% of the well) proliferating under a microscope after 5 and 10 days of culture, respectively. Cloning frequency was calculated considering the Poisson distribution (which predicts that 63% of wells should contain 1 cell following manual plating). The composition of the generated cells was evaluated on May-Grünwald/Giemsa (Merck) stained cytospin slides prepared from pooled colonies. To perform colony RT-PCRs, individual colonies were transferred to a 96-well PCR plate with PBS. After spinning down (1000g 10min), the supernatant was aspirated (leaving approximately 5μl), 5μl 2× lysis buffer (0.8% NP40, 0.125mM dNTPs, 5mM DTT and 0.005U RNaseOUT) added. Plates were frozen at −80°C till further use. Nested RT-PCRs were, with small modifications, performed as previously described (Hu et al., 1997; Adolfsson et al., 2005).

Evaluation of myeloid lineage potential and colony RT-PCRs was performed as previously described with minor modifications (Mansson et al., 2008; 2010; Hu et al., 1997; Adolfsson et al., 2005). For detail see Supplemental Materials and Methods.

### In vitro evaluation of T cell potential on OP9-DL1

Single cells were FACS sorted onto pre-seeded OP9-DL1 stromal cells in 96-well plates. OP9-DL1 co-cultures were performed in OptiMEM (with GlutaMax, Gibco) supplemented with 100U/ml penicillin-streptomycin (Hyclone), 100μM β- mercaptoethanol, 10% Fetal Calf Serum (Merck), 5ng/ml murine SCF and 5ng/ml human FL (PeproTech). Additional FL (5ng/ml) was added weekly. The clonal readout was evaluated by flow cytometry after 3 weeks of culture (for antibody panel see Table S3).

## Supporting information

Manuscript_SFigures

TableS1_RNAseq

FOXO_TableS2_ATACseq

TableS3_ab_lists

## SUPPLEMENTAL MATERIAL

Figures S1-S5

Table S1-3

## DATA AVAILABILITY

ChIPseq, ATACseq and RNAseq data from FOXO knockouts and control animals are available from the European Nucleotide Archive (ENA) under the accessions PRJEB41018 and PRJEB20316.

## AUTHOR CONTRIBUTIONS

RM conceptualized the studies. LPP, SK, NF and RM planned the study. SK, NF, MH, TB and RM performed FACS phenotyping. SK, AK, XLW and CG performed ATACseq and RNAseq. LPP, SK, NF, TB, HQ and RM performed *in vitro* colony assays. NF, ES and SL performed transplantation experiments. SK and CD performed ELISA. LPP, JH and MK analyzed omics data. ASJ, JW, NK and PW assisted with animal studies. LW, SL and RM supervised the study. LPP and RM wrote the manuscript with input from the other authors. All authors reviewed the manuscript before submission.

## ACKNOWLEDGEMENTS

We would like to thank the staff at the animal facilities (Karolinska Institutet, KI) for animal care; the Centre for Cellular Analysis (CCA, Karolinska University Hospital) for assistance with cell sorting; the MedH Flow Cytometry core facility (KI) for access to equipment; the Swedish National Infrastructure for Computing (SNIC) at Uppmax for computational resources and data storage; National Bioinformatics Infrastructure Sweden (NBIS) for providing long-term informatics support and mentoring via the Swedish Bioinformatics Advisory program; the National Genomics Infrastructure for sequencing service; Professor Joakim Dillner (KI) with colleagues for access to the Illumina NextSeq 500 system; and Assistant professor Taras Kreslavskiy (KI) for critical reading of the manuscript. This work was supported by funding from the Swedish Cancer Society (Cancerfonden), Swedish Research Council, King Gustav V Jubilee Fund (Radiumhemmet), the Swedish Foundation for Strategic Research, the Knut and Alice Wallenberg Foundation and generously, a donation by Björn and Lena Ulvaeus. In addition, the Karolinska Institutet doctoral education program (KID) supported the doctoral studies of LPP and JH. The authors have no conflicts of interest to disclose.

## ABBREVIATIONS

BCR: B cell receptor

BM: Bone marrow

CLP: Common lymphoid progenitor

CLP-A: LY6D-CLPs

CLP-B: LY6D+ B-lineage specified

CLPs CTRL: controls

DARs: Differentially accessible regions

FCS: fetal calf serum

FOXO: Forkhead box O

FOXO1ko: Conditional loss of FOXO1

FOXO3ko: Conditional loss of FOXO3

FOXOdko: Conditional loss of FOXO3 and FOXO1

Ig: Immunoglobulin

IgH: Ig heavy

IgL: Ig light

LMPP: Lymphoid-primed multipotent progenitors

PB: peripheral blood

PCA: principal component analysis

PI: propidium iodide

TFBS: transcription factor binding sites

TF: transcription factor.

## Notes

### Competing Interest Statement

The authors have declared no competing interest.

