## Supplementary material for "FOXO dictate initiation of B cell development and myeloid restriction in common lymphoid progenitors": Manuscript_SFigures

#### SUPPLEMENTAL FIGURES

### Fig. S1

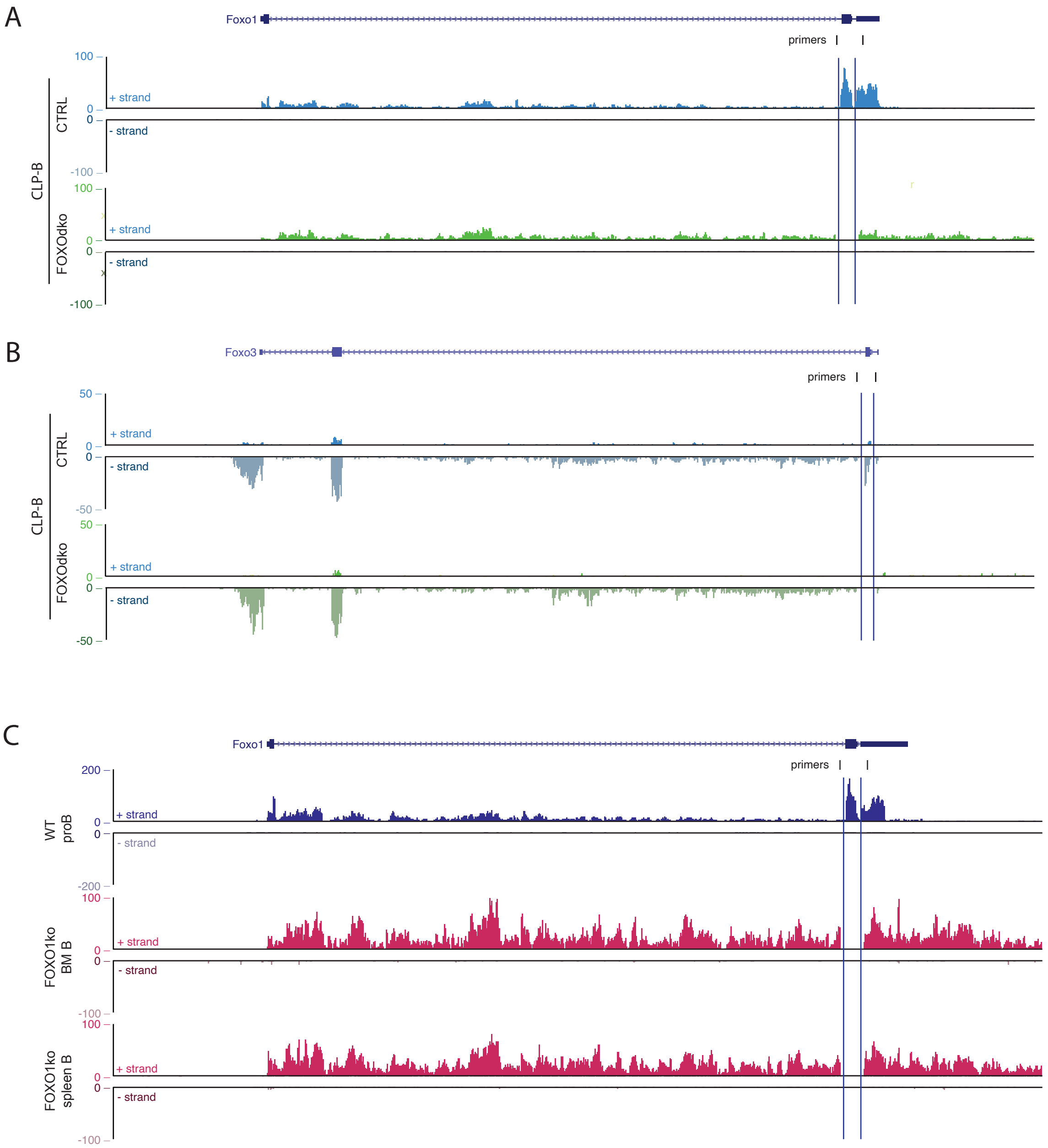
A
Foxo1
primers
100
0
-100
+ strand
- strand
CTRL
FOXOdko
CLP-B
B
Foxo3
primers
50
0
-50
+ strand
- strand
CTRL
FOXOdko
CLP-B
C
Foxo1
primers
200
0
-200
100
-100
+ strand
- strand
WT proB
FOXO1ko BM B
FOXO1ko spleen B

**Figure S1. The conditional Foxo alleles are efficiently deleted by Vav-iCre.** (A-C) Genome browser tracks showing the lack of the floxed *Foxo1* and/or *Foxo3* exons in transcripts from FOXOdko (*Foxo1*$^{ff}$*Foxo3*$^{ff}$*VAV*$^{iCre}$) CLP-Bs (A-B) and FOXO1ko B cell(C). Axis display read counts (normalized to 10 million total reads) on the plus and minus strand respectively. The position of primers flanking the floxed exons are indicated.

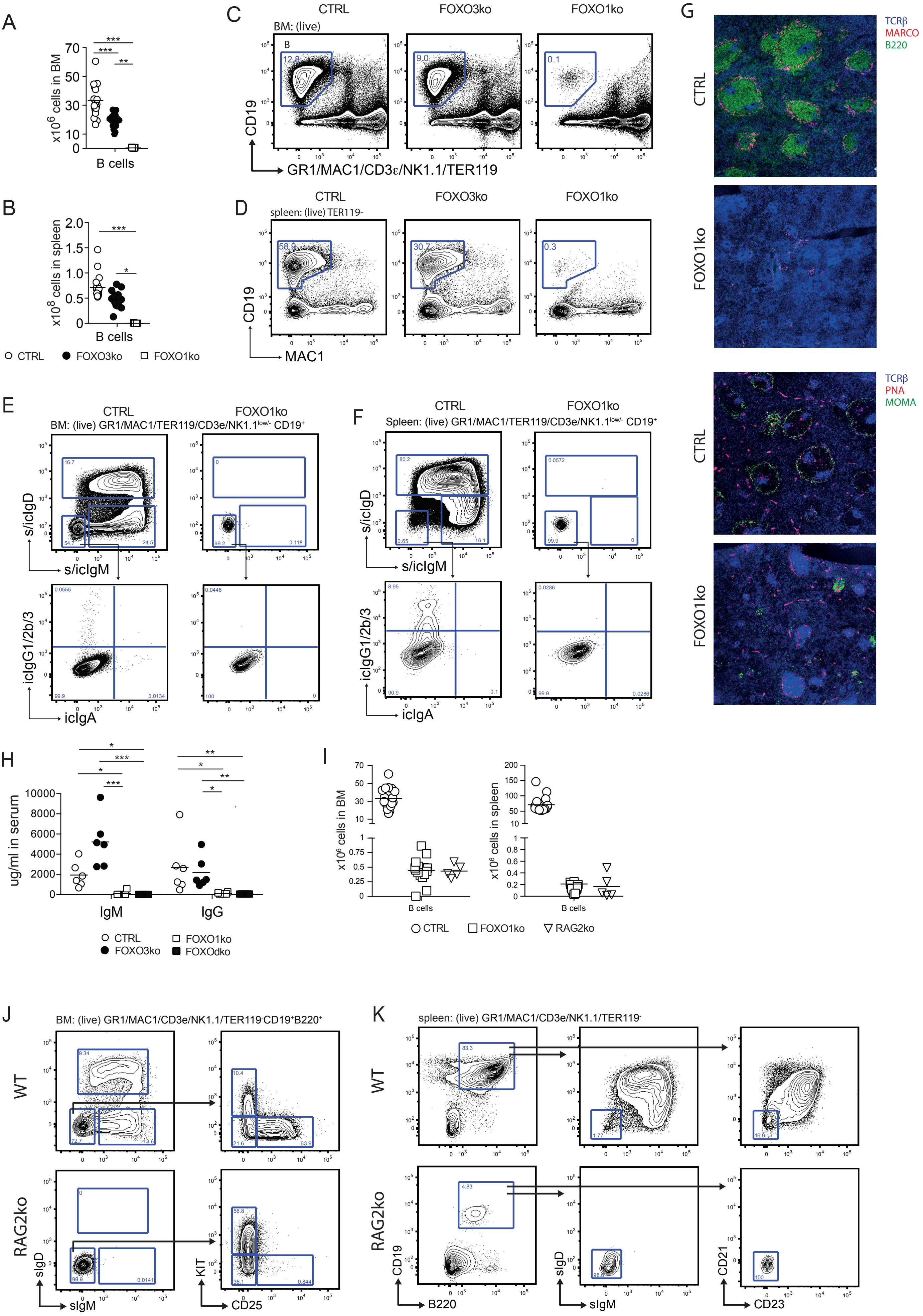
A
x10^6 cells in BM
B cells
B
x10^8 cells in spleen
B cells
CTRL FOXO3ko FOXO1ko
C
CTRL FOXO3ko FOXO1ko
BM: (live)
CD19
GR1/MAC1/CD3ε/NK1.1/TER119
D
CTRL FOXO3ko FOXO1ko
spleen: (live) TER119-
CD19
MAC1
E
CTRL FOXO1ko
BM: (live) GR1/MAC1/TER119/CD3e/NK1.1low/- CD19+
s/icIgD
s/icIgM
icIgG1/2b/3
icIgA
F
CTRL FOXO1ko
Spleen: (live) GR1/MAC1/TER119/CD3e/NK1.1low/- CD19+
s/icIgD
s/icIgM
icIgG1/2b/3
icIgA
G
CTRL
FOXO1ko
TCRβ
MARCO
B220
TCRβ
PNA
MOMA
H
ug/ml in serum
IgM IgG
CTRL FOXO1ko
FOXO3ko FOXOdko
I
x10^6 cells in BM
x10^6 cells in spleen
B cells
CTRL FOXO1ko RAG2ko
J
BM: (live) GR1/MAC1/CD3e/NK1.1/TER119-CD19+B220+
WT
RAG2ko
sIgD
sIgM
KIT
CD25
K
spleen: (live) GR1/MAC1/CD3e/NK1.1/TER119-
WT
RAG2ko
CD19
B220
sIgD
sIgM
CD21
CD23

**Figure S2. Characterization of residual B cells in FOXO and RAG knockout mice.** (A-B) Total number of B cells in (A) BM and (B) spleen. In panel A-B and H-I: each dot represents data from an individual mouse; p-values were calculated using the Kruskal Wallis test with Dunn's test of multiple comparisons; *, ** and *** indicating p-values <0.05, <0.01, and <0.001 respectively. (C-D) Gating strategy for identification of B cells in (C) BM and (D) spleen. (E-F) Representative intracellular (ic) and surface (s) staining of IgA, D, G, and M on (E) BM and (F) spleen B cells. (G) Immunohistochemistry on spleen sections. Representative areas are shown. (H) ELISA detection of IgM and IgG in serum. (I) Total number of B cells in the indicated organ. (J) Gating strategy for identification of BM B lineage cells. (K) Gating strategy for identification of spleen B cell subsets.

Fig. S3

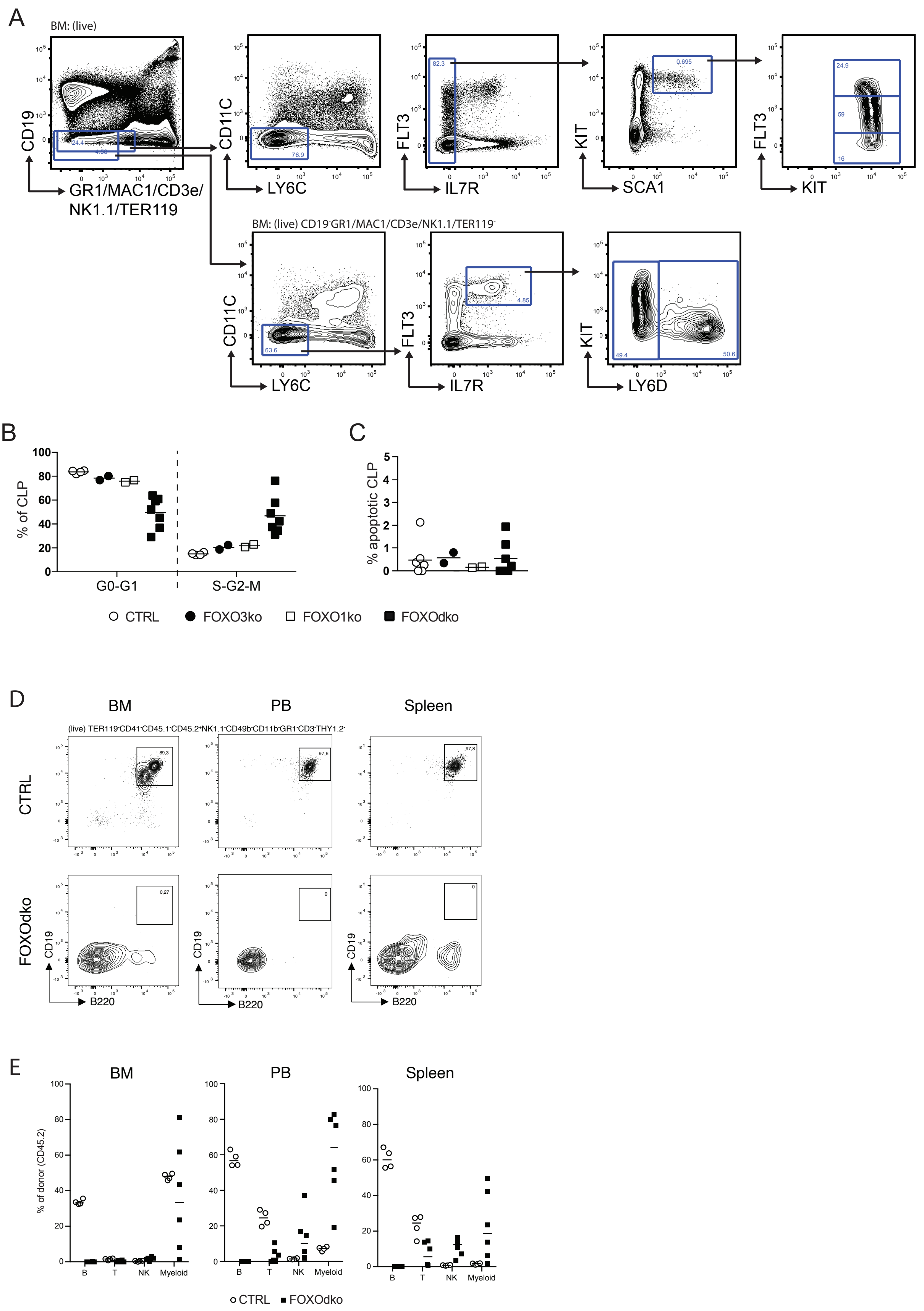
A
BM: (live)
CD19
GR1/MAC1/CD3e/
NK1.1/TER119
CD11C
LY6C
FLT3
IL7R
KIT
SCA1
FLT3
KIT
BM: (live) CD19⁻GR1/MAC1/CD3e/NK1.1/TER119⁻
CD11C
LY6C
FLT3
IL7R
KIT
LY6D
B
% of CLP
G0-G1
S-G2-M
C
% apoptotic CLP
CTRL
FOXO3ko
FOXO1ko
FOXOdko
D
BM
PB
Spleen
(live) TER119⁻CD41⁻CD45.1⁻CD45.2⁺NK1.1⁻CD49b⁻CD11b⁻GR1⁻CD3⁻THY1.2⁻
CTRL
FOXOdko
CD19
B220
E
BM
PB
Spleen
% of donor (CD45.2)
B
T
NK
Myeloid
CTRL
FOXOdko

**Figure S3. Characterization of cycling, apoptosis, and reconstitution of FOXO deficient progenitors.** (A) Gating strategy for identification of stem and progenitor cells. (B) Percentage of CLPs in the indicated cell cycle stage as determined by Ki67 and DAPI staining. (C) Percentage of apoptotic CLPs as determined by Annexin V staining. In panels B-C: each dot represents data from an individual mouse; p-values were calculated using the Kruskal Wallis test with Dunn's test of multiple comparisons; *, ** and *** indicating p-values <0.05, <0.01, and <0.001 respectively. (D) Gating strategy for identification of B cells in bone marrow (BM), peripheral blood (PB) and spleen from lethally irradiated CD45.1 mice transplanted with $5x10^6$ (CD45.2) donor (CTRL or FOXOdko) and $0.2x10^6$ support (CD45.1) BM cells. (E) Percentage of the donor ($CD45.2^+$) derived cells constituting the indicated cell-types in BM (left), PB (middle) and spleen (right). Analysis was performed 12 weeks post transplantation.

Fig. S4

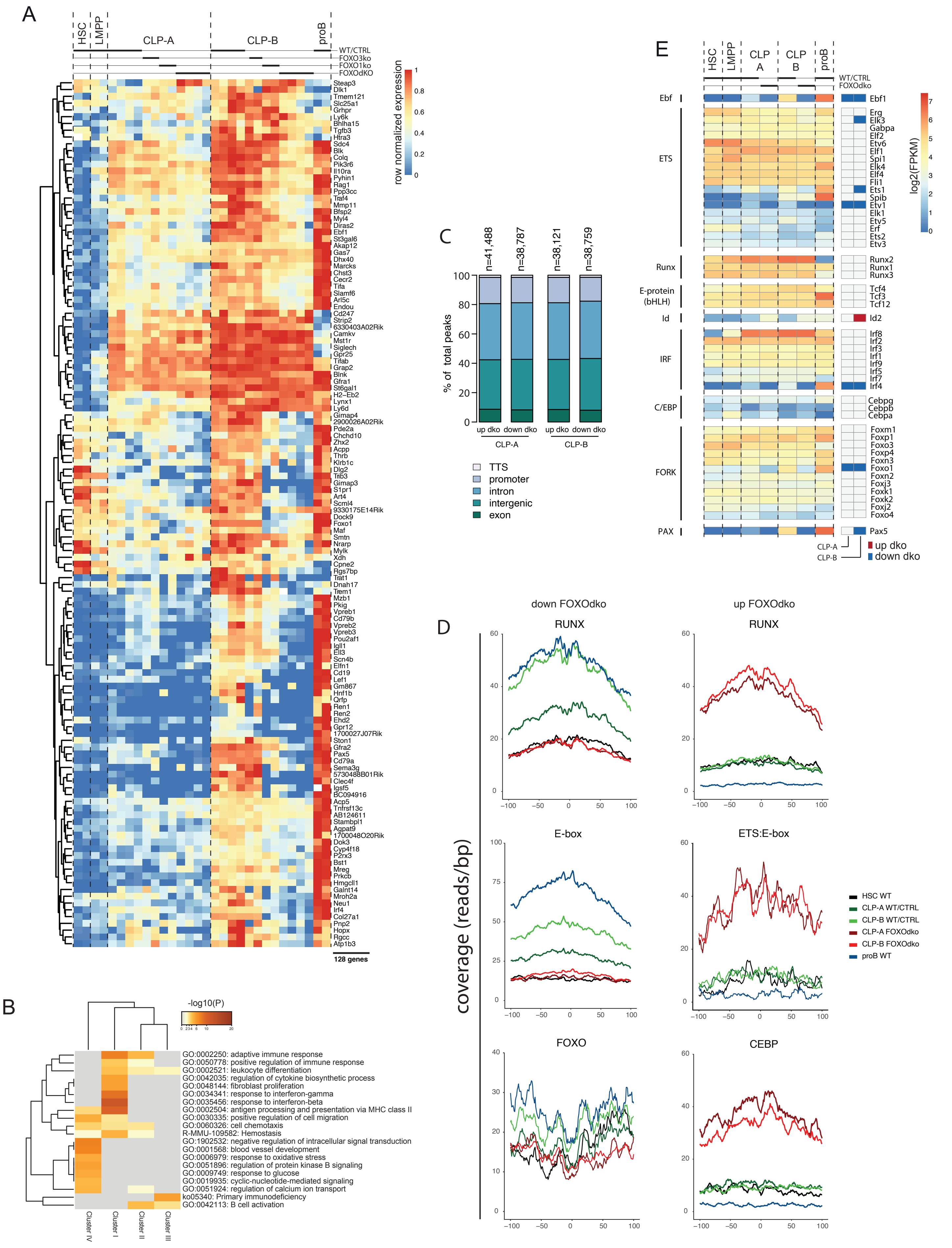
A
HSC
LMPP
CLP-A
CLP-B
proB
WT/CTRL
FOXO3ko
FOXO1ko
FOXOdKO
row normalized expression
128 genes
B
-log10(P)
GO:0002250: adaptive immune response
GO:0050778: positive regulation of immune response
GO:0002521: leukocyte differentiation
GO:0042035: regulation of cytokine biosynthetic process
GO:0048144: fibroblast proliferation
GO:0034341: response to interferon-gamma
GO:0035456: response to interferon-beta
GO:0002504: antigen processing and presentation via MHC class II
GO:0030335: positive regulation of cell migration
GO:0060326: cell chemotaxis
R-MMU-109582: Hemostasis
GO:1902532: negative regulation of intracellular signal transduction
GO:0001568: blood vessel development
GO:0006979: response to oxidative stress
GO:0051896: regulation of protein kinase B signaling
GO:0009749: response to glucose
GO:0019935: cyclic-nucleotide-mediated signaling
GO:0051924: regulation of calcium ion transport
ko05340: Primary immunodeficiency
GO:0042113: B cell activation
Cluster IV
Cluster I
Cluster II
Cluster III
C
n=41,488
n=38,787
n=38,121
n=38,759
% of total peaks
up dko
down dko
CLP-A
CLP-B
TTS
promoter
intron
intergenic
exon
D
down FOXOdko
up FOXOdko
RUNX
E-box
ETS:E-box
FOXO
CEBP
coverage (reads/bp)
HSC WT
CLP-A WT/CTRL
CLP-B WT/CTRL
CLP-A FOXOdko
CLP-B FOXOdko
proB WT
E
HSC
LMPP
CLP A
CLP B
proB
WT/CTRL
FOXOdko
Ebf
ETS
Runx
E-protein (bHLH)
Id
IRF
C/EBP
FORK
PAX
log2(FPKM)
up dko
down dko
CLP-A
CLP-B

**Figure S4. Complementary information related to gene expression and chromatin accessibility changes in FOXO deficient CLPs.** (A) Hierarchical clustering of genes differentially up-regulated (Bonferroni corrected p-value $<0.05$ and 2-fold difference and $\geq 30$ reads in at least two samples) in the CLP-A to CLP-B transition. (B) Gene set enrichment (Metascape) analysis of clusters I-IV from Figure 3D. (C) Bar graph illustrating the percentage of open chromatin regions (ATACseq peaks) within different genomic locations. The total number of peaks identified in each population is indicated above the bar-graphs. (D) Transposase integration-based cut-profiles of DARs with: decreased ATACseq signal containing RUNX, E-protein and FOXO binding sites (left); or increased ATACseq signals containing RUNX, ETS:Ebox, and CEBP binding sites (right) in FOXOdko CLPs. (E) Heatmap illustrating expression in log2(FPKM) of expressed TFs (FPKM>1) from the indicated families.

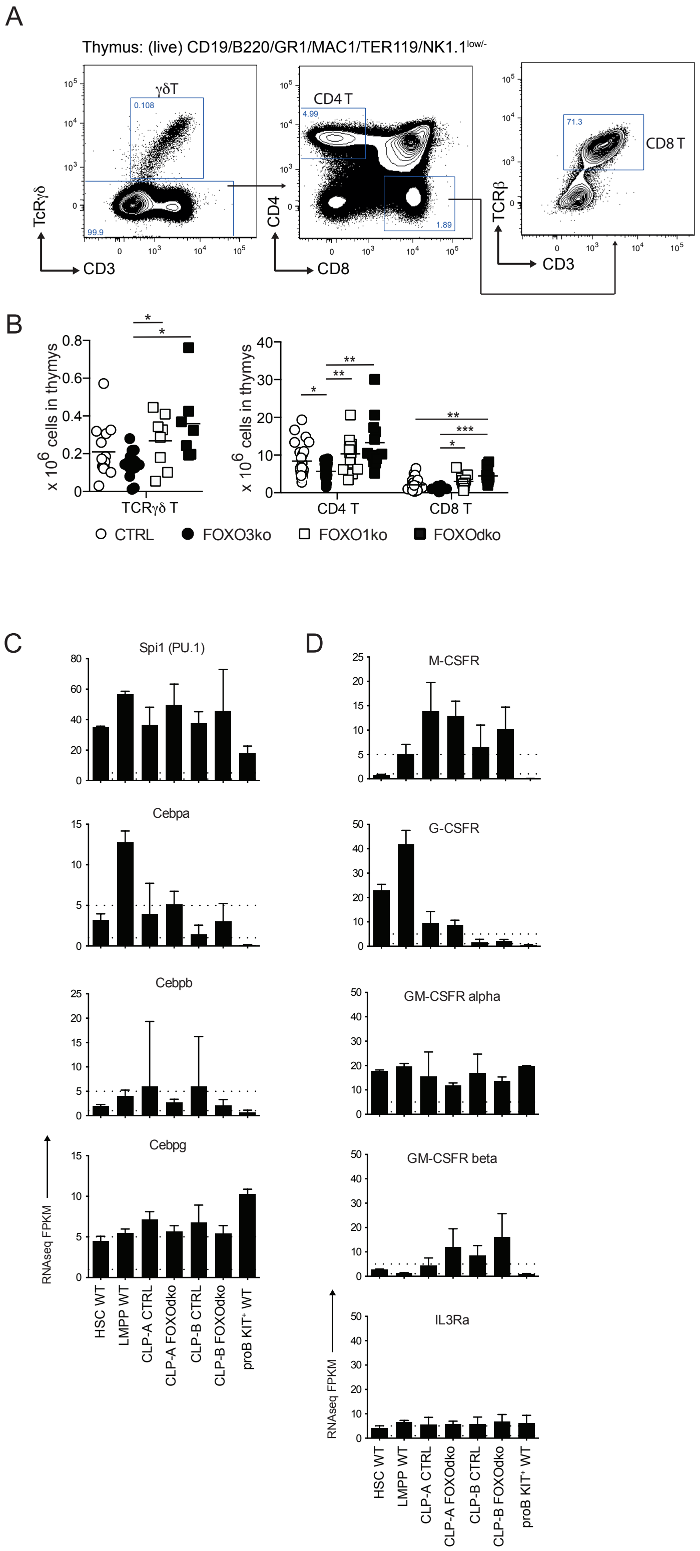
A
Thymus: (live) CD19/B220/GR1/MAC1/TER119/NK1.1low/-
γδT
CD4 T
CD8 T
TcRγδ
CD3
CD4
CD8
TCRβ
B
x 10^6 cells in thymys
TCRγδ T
CD4 T
CD8 T
CTRL
FOXO3ko
FOXO1ko
FOXOdko
C
Spi1 (PU.1)
Cebpa
Cebpb
Cebpg
RNAseq FPKM
HSC WT
LMPP WT
CLP-A CTRL
CLP-A FOXOdko
CLP-B CTRL
CLP-B FOXOdko
proB KIT+ WT
D
M-CSFR
G-CSFR
GM-CSFR alpha
GM-CSFR beta
IL3Ra

**Figure S5. Characterization of thymic populations and expression of myeloid associated genes.** (A) Gating strategy for identification of thymic populations. (B) Total number of indicated T-lineage cells. Each dot represents data from an individual mouse; p-values were calculated using the Kruskal Wallis test with Dunn's test of multiple comparisons; *, ** and *** indicating p-values <0.05, <0.01, and <0.001 respectively. (C-D) Expression (FPKM) of transcription factor (C) and cell surface receptor (D) genes associated with myeloid development.
